# The PAX3 and 7 homeodomains have evolved unique determinants that influence DNA-binding, structure and communication with the paired domain

**DOI:** 10.1101/701656

**Authors:** Gareth N. Corry, Brian D. Sykes, D. Alan Underhill

## Abstract

The PAX (<u>pa</u>ired bo<u>x</u>) family is a collection of metazoan transcription factors defined by the paired domain, which confers sequence-specific DNA-binding. Ancestral PAX proteins also contained a homeodomain, which can communicate with the paired domain to modulate DNA-binding. In the present study, we sought to identify determinants of this functional interaction using the paralogous PAX3 and 7 proteins. First, we evaluated a group of heterologous paired domains and homeodomains for the ability to bind DNA cooperatively through formation of a ternary complex (paired domain:homeodomain:DNA). This revealed that capacity for ternary complex formation was unique to the PAX3 and 7 homeodomains and therefore not simply a consequence of DNA-binding. We also found PAX3 and 7 were distinguished by an extended region of conservation N-terminal to the homeodomain (NTE). Phylogenetic analyses established the NTE was restricted to PAX3/7 orthologs of segmented metazoans, indicating it arose in a bilaterian precursor prior to separation of deuterostomes and protostomes. In DNA-binding assays, presence of the NTE caused a decrease in monomeric binding by the PAX3 homeodomain that reflected a lack of secondary structure in 1D-^1^H-NMR. Nevertheless, this inhibitory effect could be overcome by homeodomain dimerization or cooperative binding with the paired domain, establishing that protein interactions could induce homeodomain folding in the presence of the NTE. Strikingly, the PAX7 counterpart did not impair homeodomain binding, revealing inherent differences that could account for its distinct target profile in vivo. Collectively, these findings identify critical determinants of PAX3 and 7 activity, which contribute to their functional diversification.

---

The PAX family of transcription factors is defined by the <u>pa</u>ired bo<u>x</u>, a DNA binding domain first described in the *Drosophila* protein paired (Prd) (1-3). The Prd protein also contains a homeodomain and phylogenetic studies established these features characterized the ancestral state, reflecting a domain-capturing event during early metazoan evolution (4-7). In mammals, a combination of gene duplication and domain loss has given rise to a complement of nine *PAX* genes. These are assigned to 4 paralogous groups: PAX1/9, PAX2/5/8, PAX3/7, and PAX4/6 (2,3,8). Within this scheme, PAX1 and 9 have undergone complete homeodomain loss, while the PAX2, 5, and 8 proteins contain only an N-terminal portion (2,3,8). Although it is clear these primary structure differences can alter DNA-binding capacity between groups (9), how functional diversification is achieved amongst paralogs is not well understood.

The availability of disease-causing mutations in PAX proteins has provided key reagents for their biochemical characterization. Notably, for PAX3 and PAX6, mutations in either DNA-binding domain were found to affect the other, establishing that the paired domain and homeodomain can functionally interact (10-15). This behavior is also consistent with optimal binding sites for Prd where the two domains can bind cooperatively to juxtaposed recognition elements even when expressed as separate polypeptides (16,17). In the case of PAX3, a further consequence of the functional interaction is that the DNA-binding properties of the homeodomain are altered (15). This also extends to its subnuclear localization in cells, which undergoes a marked change upon tethering to the paired domain (18). It is therefore apparent intramolecular interactions can play a significant role in shaping the activity of individual PAX proteins.

In the present study, we have assessed how communication between the paired domain and homeodomain influences their overall activity, with a focus on PAX3 and PAX7. Using a variety of approaches, we found that key determinants for the functional interaction were conferred by the homeodomain. Specifically, the ability to cooperatively bind DNA in trans was universal across paired domain paralogs, but only possessed by a subset of homeodomains (PAX3 and PAX7). The PAX3 and 7 homeodomains were further distinguished by a conserved amino-terminal extension (NTE). In addition to establishing the phylogenetic history of this sequence, we found that the NTE contributes to inhibition of homeodomain DNA-binding and folding for PAX3 but not PAX7. Inhibition was overcome by homeodomain dimerization or binding with the paired domain. Together, the data support a model in which the NTE has evolved to differentially regulate homeodomain activity and functional interaction with the paired domain for closely related PAX proteins. Our findings also emphasize the critical importance of sequences outside of the canonical DNA-binding domains in modulating nucleic acid interactions.

## EXPERIMENTAL PROCEDURES

### Bacterial protein expression and electrophoretic mobility shift assays

Expression and purification of recombinant proteins and electrophoretic mobility shift assays (EMSAs) were carried out as previously described (19). Briefly, all expression plasmids were created in pET21a (Novagen; N-terminal T7•tag® and C-terminal His•tag®) and proteins were purified by nickel chromatography carried out in batch format. A full list of recombinant proteins including amino acid boundaries and corresponding oligonucleotide primers is provided in Table 1. Phospho-site mutations (corresponding to codons for Ser-201, 205, and 209 in PAX3) were made using the PAX3 196-279 backbone and substitution to alanine or aspartic acid codons by PCR. Template cDNAs were described in previous publications (15,18,19) or purchased from Open Biosystems (Dharmacon/GE Life Sciences) as part of the Mammalian Gene Collection.

### Phylogenetic analyses

The human PAX3 protein (NCBI Reference Sequence: NP_852122.1) was used as an entry point to collect homologous sequences using a standard protein BLAST search. Additional searches that focused on specific phyla and subkingdoms were carried out by standard protein BLAST or TBLASTN to search multiple translated nucleotide databases, which provided access to a large number of PAX homologues not currently annotated. Collectively, these searches broadly covered Parazoa, Mesozoa, and Eumetazoa. Within the latter phylum, more extensive searches were carried out within Protostomia and Deuterostomia descendants to ensure adequate coverage. Sequences were aligned with Clustal Omega (http://www.ebi.ac.uk/Tools/msa/clustalo/) and depicted with BoxShade (http://www.ch.embnet.org/software/BOX_form.html). Species key is as follows: cbri, *Caenorhabditis briggsae*; crem, *Caenorhabditis remanei*; cele, *Caenorhabditis elegans*; asum, *Ascaris suum*; ct, *Capitella teleta*; ap, *Alvinella pompejana*; he, *Helobdella robusta*; cin, *Ciona intestinalis*; hr, *Halocynthia roretzi*; hsa/hsap, *Homo sapiens*; ggal, *Gallus gallus*; drer, *Danio rerio*; bbe, *Branchiostoma belcheri*; bf, *Branchiostoma floridae*; ebu, *Eptatretus burgeri*; ami, *Apis mellifera*; cflor, *Camponotus floridanus*; hsalt, *Harpegnathos saltator*; api, *Acyrthosiphon pisum*; phum, *Pediculus humanus corporis*; tcast, *Tribolium castaneum*; same, *Schistocerca Americana*; dmel, *Drosophila melanogaster*; ptepi, *Parasteatoda tepidariorum*; cgig, *Crassostrea gigas*; obi, *Octopus bimaculoides*; nve, *Nematostella vectensis*; amil, *Acropora millepora*; aja, *Anthopleura japonica*; tad, *Trichoplax adhaerens*.

### Nuclear magnetic resonance spectroscopy

Folding of bacterially expressed PAX3 proteins comprising the homeodomain with (196-279) and without (219-279) the N-terminal extension was assessed by 1D ^1^H NMR spectroscopy. Using conditions we have previously established for soluble purification of PAX3 by Ni^2+^-agarose chromatography (19), protein was recovered and concentrated to 4 mg/mL prior to dialysis into 100 mM NaCl/0.5 mM Imidazole. ^1^H NMR spectra were obtained using an 800 MHz spectrometer available through the NANUC (National High Field Nuclear Magnetic Resonance Centre) facility at the University of Alberta with an acquisition time of 2 s and 384 scans per spectrum.

## RESULTS

### Determinants of paired domain-homeodomain functional interaction

Of the nine mammalian PAX members, PAX3 and PAX7 form an evolutionary subgroup that includes *Drosophila* Prd and gooseberry (Gsb) (8). These proteins can engage DNA using various combinations of the paired domain and homeodomain, leading to a range of distinct target sites (3,8). For Prd, these have been empirically derived by in vitro selection, which identified 3 primary modes of binding (17). Of these, the highest affinity coincided with binding of the paired domain and homeodomain to juxtaposed recognition elements, and was also supported by in vivo analyses (16,17). Moreover, the Prd paired domain and homeodomain could bind synergistically to these elements when expressed as separate polypeptides to form a ternary complex (17). In this context, we have previously shown that the PAX3 paired domain and homeodomain can cooperate when binding as individual proteins to a regulatory element from the *MITF* promoter, although each bound poorly in isolation (19). We therefore exploited this attribute to evaluate the ability of heterologous domain combinations to cooperate in DNA-binding.

To determine the specificity of the functional interaction, we evaluated the ability of the PAX3 paired domain or homeodomain to form a ternary complex with counterparts from other PAX proteins or the broader homeodomain superfamily (Figure 1). Overall sequence identity with PAX3 ranged from 93% (PAX7) to 66% (PAX6) within the paired domain (Figure 1A), and 95% (PAX7) to 42% (MSX1) within the homeodomain, dropping to 28% for the partial domain in PAX2 (Figure 1B). Although paired domains from all 4 paralogous groups (PAX1, 2, 6, and 7) could form a ternary complex with the PAX3 homeodomain (Figure 1C), this was not reciprocal. Specifically, only the paralogous PAX7 homeodomain could form a ternary complex with the PAX3 paired domain (Figure 1D). Neither the PAX6 homeodomain, nor the partial version from PAX2 where able to do so and this was also seen with PITX2, PRRX1, and MSX1 (Figure 1D), all of which belong to the same Prd evolutionary subclass. Notably, the ability to form a ternary complex amongst paired domains did not correlate with overall sequence identity, given the PAX2 paired domain was most efficient, but was generally related to the level of paired domain DNA binding (Figure 1C). These data indicate that the key determinants for cooperative binding reside within the homeodomain.

**FIGURE 1.**
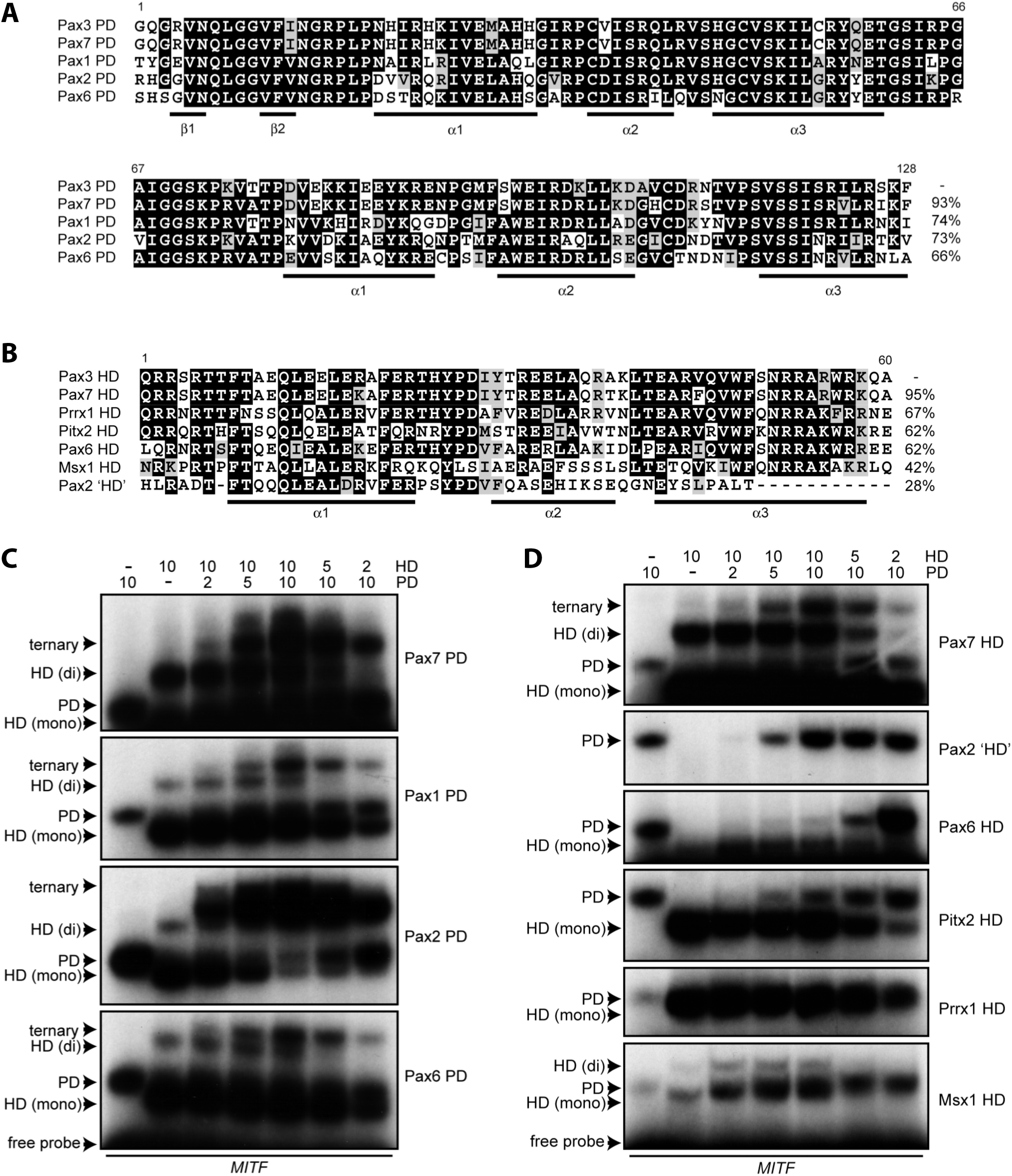
Ternary complex formation amongst heterologous paired domains and homeodomains. (a) Sequence alignment of paired domains with corresponding amino acid identity indicated (identical resides are indicated by *black* highlighting). (b) Sequence alignment of prd-type homeodomains with corresponding amino acid identity indicated (identical resides are indicated by *black* highlighting). In both a) and b), secondary structure features are indicated below the sequence. (c) Ternary complex assays using heterologous paired domain and PAX3 homeodomain combinations in electrophoretic mobility shift assays. Relative amounts of paired domain and homeodomain are indicated numerically. Individual complexes are denoted as ternary (PD:HD:DNA), homeodomain monomer (HD (mono)), homeodomain dimer (HD (dimer)), or paired domain (PD). (d) Ternary complex assays using heterologous homeodomain and PAX3 paired domain combinations in electrophoretic mobility shift assays. Labeling as in panel (c). EMSAs are representative of a minimum of 2 independent pilot experiments.

### Identifying specificity determinants within the PAX3 and PAX7 homeodomains

Using a chimera approach, we sought to narrow the search for homeodomain determinants by interchanging helical segments between PAX3, PRRX1 and PITX2. Nevertheless, chimeric proteins exhibited no (PRRX1/PAX3) or greatly reduced (PITX2/PAX3) DNA binding (not shown), suggesting the basis for their distinct activities could not be mapped to a discrete segment of the homeodomain. Together with the results from ternary complex assays (Figure 1), it appears the PAX3 and PAX7 homeodomains have evolved features that influence their structure and communication with the paired domain. We therefore evaluated sequence features across the PAX and broader homeodomain superfamily, which revealed an 18 amino acid region of homology contiguous with the homeodomain N-terminus that was unique to the PAX3/PAX7/Prd/Gsbn subgroup (Figure 2A). A WebLogo depiction of the motif shows that the most conserved features are the S-D/E pairs at positions 205 and 209, although PAX3 orthologs contain a third pair at position 201 (Figure 2B). Moreover, phylogenetic analyses established that this N-terminal extension (NTE) was further restricted to PAX3/PAX7/Prd/Gsbn proteins from segmented metazoans. Together with its absence in ancestral PaxB predecessors, the NTE likely arose in a common bilaterian precursor and we propose that it should be considered as part of the homeodomain.

**FIGURE 2.**
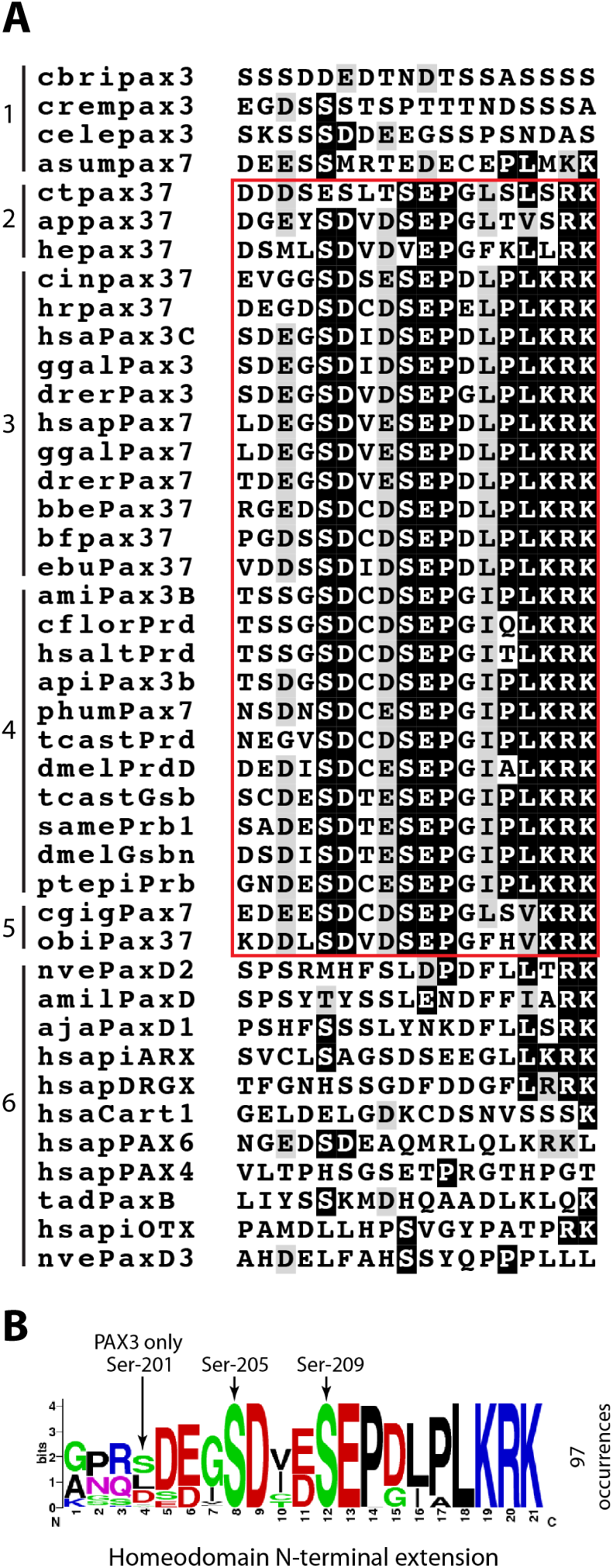
The PAX3/PAX7/Prd/Gsbn proteins contain a highly conserved homeodomain amino-terminal extension. (a) Sequence alignment of the 18 amino acid region preceding the homeodomain (identity depicted as *white on black*; conservative changes in *gray*). The NTE (*red box*) appears in the PAX3/PAX7/Prd/Gsbn subgroup from annelids (*2*), chordates (*3*), arthropods (*4*) and mollusks (*5*). The motif is absent from nematode (*1*), ancestral PAX proteins and related homeodomains (*6*). (b) WebLogo depiction of the NTE derived from analysis of 97 distinct occurrences of PAX3/7 orthologs. Conserved serine residues are indicated at positions 201, 205 and 209.

### The NTE affects PAX3 homeodomain DNA-binding and folding

Electrophoretic mobility shift assays (EMSAs) were carried out to evaluate DNA-binding by the PAX3 homeodomain in the presence of the NTE and varying portions of the 57 amino acid linker to the paired domain (Figure 3A). Oligonucleotides assessed monomeric (*P1/2*) or dimeric (*P2*) homeodomain binding (20) and cooperative binding with the paired domain (*MITF*). Significantly, the presence of the NTE (196-279) caused a reduction in monomeric binding (Figure 3B, *P1/2*, lane 3) compared to the canonical homeodomain (Figure 3B, *P1/2*, lane 5) or a version containing the nuclear localization sequence (Figure 3B, *P1/2*, lane 4). Although a further reduction occurred with the full linker (Figure 3B, *P1/2*, lane 1), these results establish that the NTE modulates homeodomain activity. In contrast, all proteins efficiently bound the dimeric *P2* site (Figure 3B, *P2* panel) and, with the exception of the full linker derivative, all were effective at ternary complex formation (Figure 3B, *MITF;* compare lane 1 to lanes 2-5). Given these effects, folding was assessed for the homeodomain and its NTE counterpart using 1D ^1^H NMR spectroscopy (Figure 4). The spectral peaks indicated the PAX3 homeodomain was folded in the absence of DNA (Figure 4, 219-279), but their loss in the NTE derivative (Figure 4, 196-279) signified a lack of structure. The NTE therefore renders the homeodomain refractory to folding when binding as a monomer (Figure 3B, *P1/2*, lane 3) but can be restored by homeodomain-homeodomain or paired domain-homeodomain interaction (Figure 3B, *P2* and *MITF*). This suggests cooperative binding can induce a disorder to order transition that overcomes the inhibitory effect of the NTE.

**FIGURE 3.**
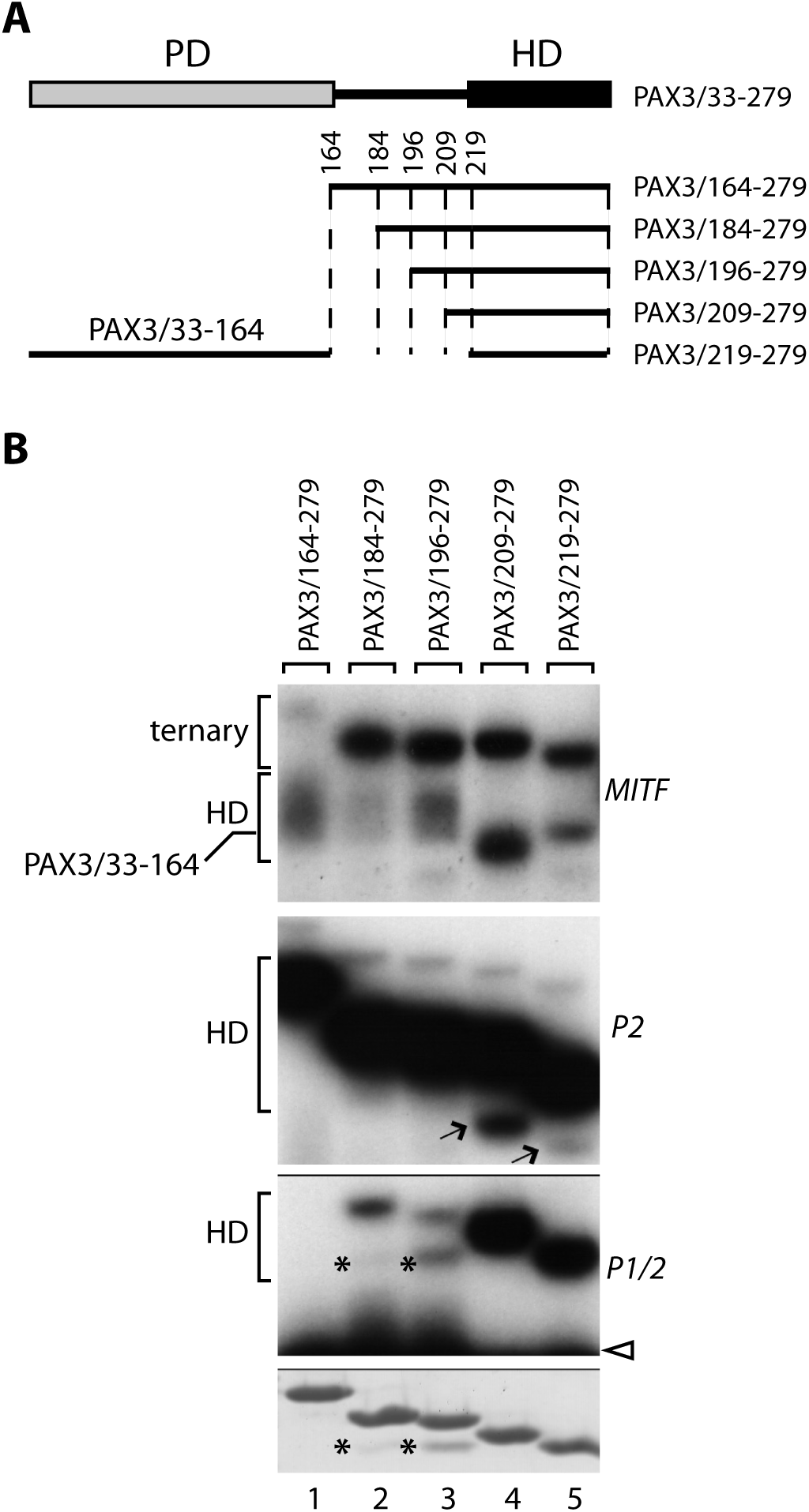
The NTE modulates DNA-binding by the PAX3 homeodomain. (a) Schematic illustrating the boundaries for recombinant PAX3 proteins used, which span various portions of the paired domain (PD), homeodomain (HD) and linker region. The carboxy terminal segment of PAX3 is not shown. (b) Recombinant proteins comprising the core homeodomain (219-279) and increasing portions of the linker (209-279, 196-279, 184-279 and 164-279) were used in EMSAs to test monomeric (*P1/2*) and dimeric (*P2*) homeodomain binding, and ternary complex formation with the paired domain (*MITF*). In the *MITF* panel, location of the paired domain (*PD*), homeodomain (*HD*) and ternary (*PD/HD*) complexes are indicated. Unbound probe (*open arrowhead*) is shown for *P1/2*. A coomassie gel indicates equal loading in the lower panel. The *asterisk* denotes a minor proteolytic fragment. EMSAs are representative of a minimum of 2 independent pilot experiments.

**FIGURE 4.**
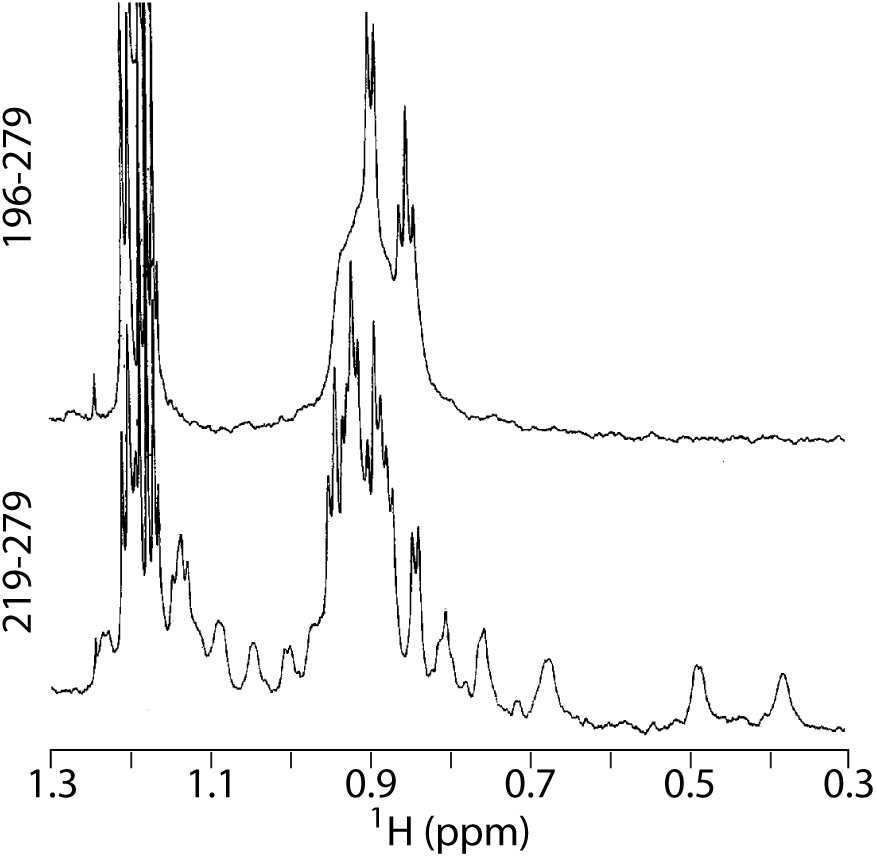
The NTE negatively affects folding of the PAX3 homeodomain. One-dimensional proton (^1^H) NMR spectroscopy of the core homeodomain (219-279) and NTE derivative (196-279). Spectra show ^1^H resonances in parts per million (ppm).

### The NTE differentially affects DNA-binding by PAX3 and PAX7

Despite extensive sequence identity between PAX3 and PAX7, they exhibit distinct genomic distributions in vivo (21). Given this, we assessed how the NTE influenced DNA-binding by the PAX7 homeodomain. Strikingly, although the core homeodomain for PAX3 and PAX7 exhibited identical binding to *P1/2* and *P2*, the NTE did not diminish PAX7 monomeric binding (Figure 5A, *P1/2*, compare lanes 1-5 to 6-10). Likewise, only the PAX7 NTE-HD exhibited appreciable levels of monomeric complex on *P2* (Figure 5A, *P2*, lanes 6 and 7). These data support a prominent role for the NTE in control of DNA-binding by paralogous homeodomains. At the same time, we found the isolated paired domain of PAX7 displayed lower affinity than PAX3 for its optimal binding site (Figure 5B). Importantly, these differences are consistent with ChIP-seq analyses using full-length proteins where PAX7 displayed a marked preference for homeodomain sites, while PAX3 bound mostly via the paired domain (21). We therefore evaluated how contiguous polypeptides spanning the paired domain and homeodomain of PAX3 and PAX7 interacted with binding sites for each domain (Figure 5C). This recapitulated the ChIP-seq findings (21) where the PAX3 PDHD protein exhibited a preference for the optimal paired domain motif (Figure 5C, *Nf3’*), while the PAX7 version displayed elevated binding to the monomeric homeodomain site (Figure 5C, *P1/2*, compare lanes 2 and 3 to 7 and 8). The PDHD PAX3 protein does however ameliorate the effect of the NTE to partially restore *P1/2* binding. Consequently, the distinct activities observed for the isolated PAX3 and 7 domains carry over to the tethered derivatives and reflect their in vivo binding preferences. Specifically, it appears reduced binding by the PAX7 paired domain together with elevated homeodomain binding can account for the different profile in vivo.

**FIGURE 5.**
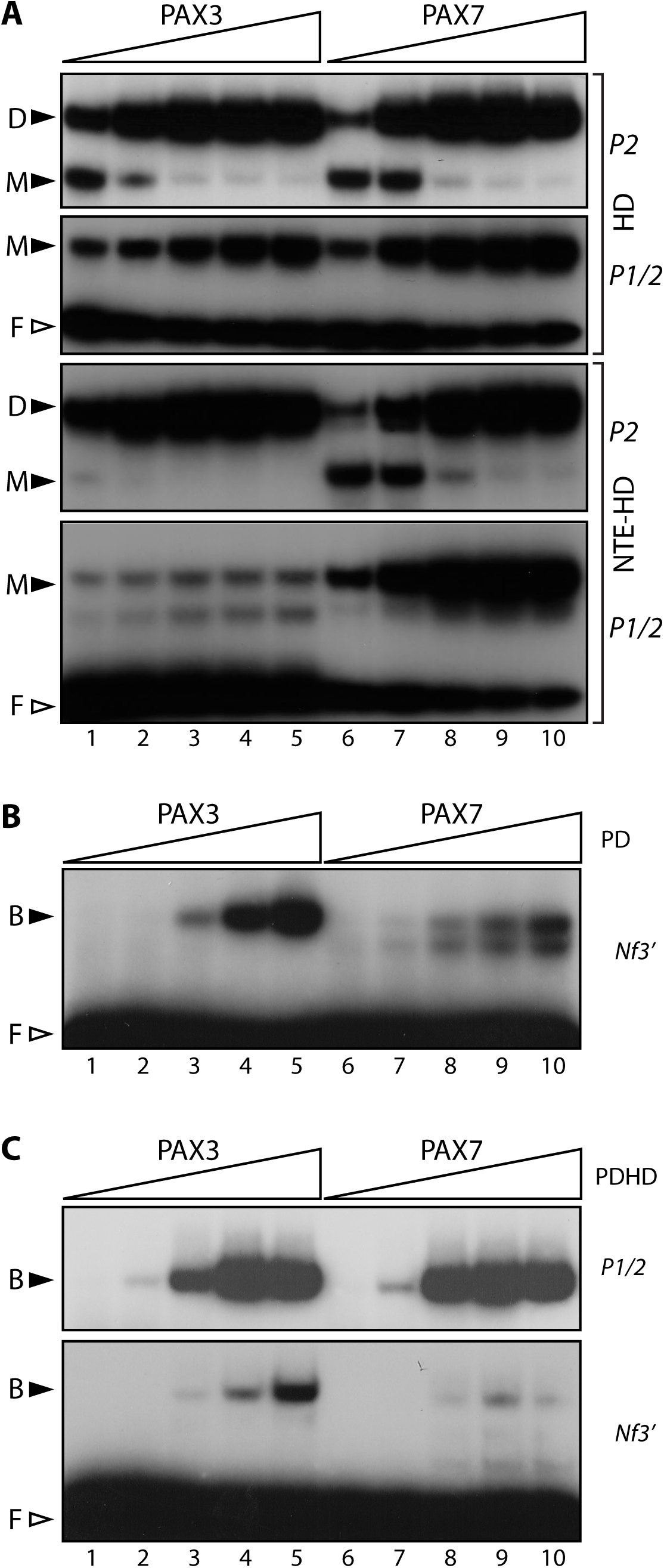
Differential effect of the NTE on DNA-binding by the PAX3 and PAX7 homeodomains. (a) EMSAs compare monomeric (*P1/2*) and dimeric (*P2*) binding of the core PAX3 and PAX7 homeodomains (top 2 panels) to their NTE counterparts (lower 2 panels). Protein levels increase from left to right. Unbound probe (*F, open arrowhead*) is shown for *P1/2* and the location of monomer (*M*) and dimer (*D*) complexes is indicated. (b) EMSAs compare binding of the isolated PAX3 and PAX7 paired domains to an optimal binding site (*Nf3’*). Protein levels increase from left to right. Unbound probe (*F, open arrowhead*) is shown for *Nf3’* and the location of bound (*B*) complexes is indicated. (c) EMSAs compare binding of PAX3 and 7 polypeptides spanning the paired domain and homeodomain (PDHD) to assess monomeric homeodomain (*P1/2*) or paired domain (*Nf3’*) binding. Unbound probe (*F, open arrowhead*) is shown for *Nf3’* and the location of bound (*B*) complexes is indicated. EMSAs are representative of a minimum of 2 independent pilot experiments.

### Phosphorylation within the NTE has the potential to modulate PAX3 DNA-binding

The most conserved features of the NTE (excluding the Lys-Arg-Lys portion of the nuclear localization signal) are the Ser-Asp/Glu pairs at positions 205 and 209 (Figure 2B). Amongst PAX3 orthologs, Ser-201 is also highly conserved and followed by an acidic residue (Figure 2B). These 3 serine residues are known to undergo phosphorylation by Casein Kinase-1/2 (CK1/2) and Glycogen Synthase Kinase-3ß (GSK3ß) (22-26). To test the influence of phosphorylation on DNA-binding, we used phosphomimetic mutants that replace the 3 serine residues in PAX3 with aspartic acid, together with alanine substituted controls. These were compared to wt PAX3/196-279, as well as derivatives lacking the NTE (209-279 and 219-279), in EMSAs that assessed *P2* and *P1/2* binding, and ternary complex formation (as in Figure 3). Intriguingly, the Ser→Ala mutants recovered monomeric DNA-binding (Figure 6A, *P1/2*, compare lanes 3 and 4), while Ser→Asp showed an intermediate effect (Figure 6A, *P1/2*, compare lanes 3 and 5; the triple-Asp mutant has increased mobility on *P1/2* and *P2*). This indicates that the serine residues contribute to the inhibitory effect of the NTE on monomeric binding. The Ser→Asp mutant differed by not fully recovering upon dimerization to *P2* (Figure 6A, *P2*, compare lanes 3-5) and not forming a ternary complex on *MITF* (Figure 6A, *MITF*, lane 5). Although the wt and Ser→Ala mutant appear similar in ternary complex formation, differences appeared when assessed over a range of protein concentrations (Figure 6B). In particular, the Ser→Ala mutant was more efficient at ternary complex formation when the homeodomain was limiting, whereas the wt protein formed a complex with increased mobility (Figure 6B, lanes 6 and 7, compare *WT* to *AAA*). Although this complex is absent from the homeodomain control (Figure 6B, lane 2), its position corresponds with that of a homeodomain dimer and would imply that the paired domain can influence binding in *trans*. Under the same conditions, the Ser→Asp mutant does not show appreciable binding on its own or with the paired domain (Figure 6B, *DDD*). Collectively, the NMR and DNA-binding analyses establish that the serine residues (or a subset therein) confer an unfolded ground state that inhibits monomeric binding, and that phosphorylation has the potential to affect the innate properties of the homeodomain and communication with the paired domain.

**FIGURE 6.**
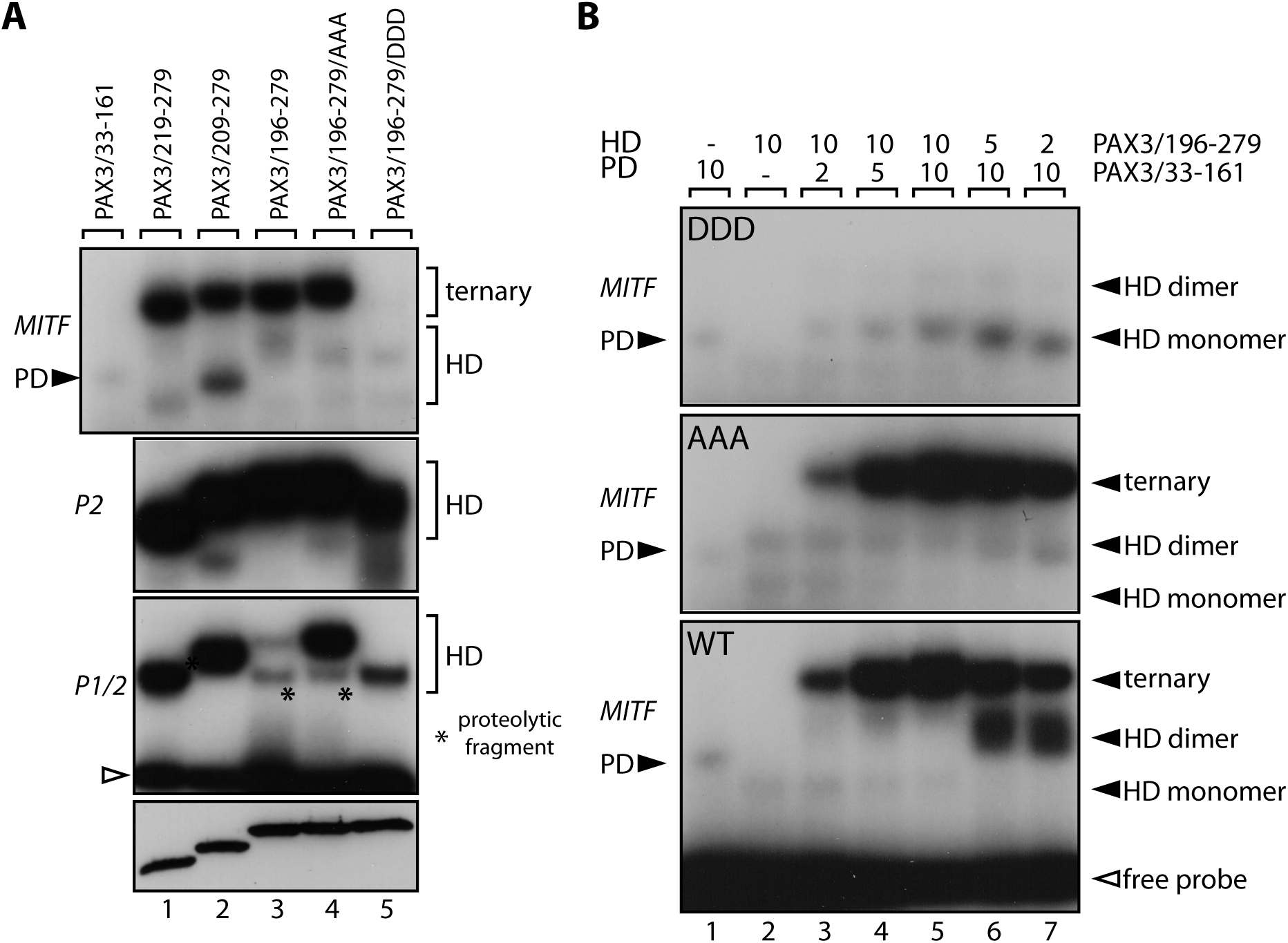
Modification of NTE serine residues affects homeodomain binding and communication with the paired domain. (a) EMSAs were carried with indicated proteins (see Figure 3A for schematic). Substitutions of Serine-201/205/209 were made with Alanine (196-279/AAA) and the phosphomimetic Aspartic acid (196-279/DDD). In the *MITF* panel, a paired domain-only control is included (33-164). The lower panel indicates equal loading by immunoblotting. The *asterisk* denotes a minor proteolytic fragment. EMSAs test monomeric (*P1/2*) and dimeric (*P2*) homeodomain (*HD*) binding, and ternary complex formation (*MITF*) with the paired domain (*PD*). In the each panel, location of the paired domain (*PD*), homeodomain (*HD*) or ternary complexes are indicated. Unbound probe (*open arrowhead*) is shown for *P1/2*. (b) Ternary complex formation (*MITF*) was tested in EMSAs using the wild type (*WT*) and Alanine (*AAA*) or Aspartic acid (*DDD*) mutants over a range of protein concentrations (relative amounts indicated numerically). Location of paired domain (*PD*), homeodomain monomer (*M*) and dimer (*D*), and ternary complexes are indicated (*closed arrowheads*). The location of the free probe (*open arrowhead*) is shown for the bottom panel. EMSAs are representative of a minimum of 2 independent pilot experiments.

## DISCUSSION

Mammalian PAX proteins are assigned to 4 paralogous groups that differ in domain content. Two of these groups, comprising PAX3/7 and PAX4/6, retained the ancestral state that was characterized by having both the paired domain and homeodomain (4,5,7,8). Functional interaction between these domains has been demonstrated for the most part using PAX3 (10,11,14,15,18,19,27-31), but is also supported with analyses of PAX6 (12,13). As a result, the two domains may have initially cooperated in trans, thereby creating selective pressure for the homeodomain capturing event (4,5). This model would be consistent with the fact that paired domains from all 4 subgroups could cooperate with the PAX3 homeodomain in ternary complex formation (Figure 1C) and with combinatorial analyses of multiple *Drosophila* and mammalian paired and homeodomain proteins (17). Nevertheless, in contrast to the study of Jun et al (17), we observed pronounced selectivity of the PAX3 paired domain, which may reflect two phenomena. First, that the PAX3 paired domain has evolved unique specificity requirements that are only satisfied by the PAX3 and 7 homeodomains. Second, that the *MITF* site has suboptimal recognition motifs for both domains (19) and requires a higher level of cooperativity than conveyed by heterologous homeodomains. Collectively, these data suggest that PAX proteins exhibit differential communication between the paired domain and homeodomain that can play an important role in target site selection.

Although the PAX version of the homeodomain (Prd type) is characterized by the canonical helix-turn-helix structure, it is distinguished by formation of a dimer that is stabilized by intermolecular interactions (20,32,33). When this is considered in the context of the NTE, it is noteworthy that dimerization was required to overcome the unfolded ground state of the PAX3 homeodomain (Figures 3B and 4). Likewise, homeodomain binding could also be induced by complex formation with the paired domain (Figure 3B), which is consistent with analyses of chimeric PAX3 proteins (11). In particular, replacement of a region corresponding to the NTE with a segment from a related Prd-class protein abrogated homeodomain DNA-binding. Consequently, using both isolated domains and intact proteins, these findings establish that that NTE is a critical regulator of homeodomain activity and communication with the paired domain. This adds to previous findings where the PAX3 paired domain altered DNA-binding, structure and subnuclear localization of the homeodomain (11,15,18,27,29). Together, it appears that the PAX3 homeodomain is constrained under normal circumstances and extensively modulated by inter and intramolecular interactions.

In the present study, we found that the PAX3 and 7 homeodomains share the ability to cooperate with the paired domain in DNA-binding (Figure 1), but acquired markedly distinct activities in the presence of the NTE (Figure 5). This is not inconsistent with their roles in vivo, which include distinct and overlapping functions during development and postnatally (34-36). For instance, PAX7 cannot rescue all facets of PAX3 function when knocked into the *Pax3* locus in mice (35), indicating inherent differences in protein activity. This idea is also supported by ChIP-seq analysis in myoblasts where PAX3 displayed a preference for paired domain motifs, while PAX7 bound primarily via the homeodomain (21). Our results support a prominent role for the NTE, together with altered paired domain binding activity, in conferring these differences (Figure 5), which were further reflected in binding profiles for the PDHD versions of these proteins. Moreover, phosphorylation of the NTE (discussed below) may create functionally distinct PAX3 and PAX7 populations, as well as those that account for their common activities through convergence of specificity.

The most conserved feature of the NTE is the serine residues at positions 205 and 209 (PAX3 numbering), which in each case are followed by an acidic residue (Figure 2B). For PAX3, a third, highly conserved S-D/E pair precedes at position 201. These serine residues are thought to constitute the only phosphorylation sites in PAX3 and are targets of GSK3β and CK1/2 (23-26). Moreover, PAX3 phosphorylation is altered in disease states, establishing these modifications as important determinants of normal and pathogenic PAX3 activity (37,38). From a biochemical standpoint, the serine residues had a two-pronged effect (Figure 6). First, they conferred a repressive effect on homeodomain binding, which was ablated by substitution with alanine. Second, introduction of a phosphomimetic residue partially restored homeodomain DNA-binding, but abrogated interaction with the paired domain. Importantly, these findings reinforce that the NTE can modulate homeodomain DNA-binding and communication with the paired domain. Mechanistically, the effect of the NTE on homeodomain folding and its control by phosphorylation are concordant with a model involving regulated order/disorder transitions of an autoinhibitory domain (39-45). In support of this, substitutions at Ser-201, 205 and 209 reduce the disorder probability using DisEMB™ (46). This model would provide an important means for upstream signaling pathways to fine tune target gene selection and regulatory outputs for individual PAX proteins by modulating domain use.

Phylogenetic analyses established that the NTE was contiguous with the homeodomain in all occurrences, suggesting they are under spatial constraint and effectively redefining the domain boundary. Moreover, the fact that the NTE was restricted to the PAX3/PAX7/Prd/Gsbn subclass meant it invariably occurred in conjunction with the paired domain. We therefore propose that these three entities have co-evolved through extensive functional interplay within this PAX subgroup, which is highlighted by the lack of a corresponding event amongst PAX4 and 6 paralogs. Intriguingly, the NTE was further limited to segmented metazoans, being notably absent from nematode Pax37 orthologs (Figure 2A). This phylogenetic distribution suggests the NTE responds to a segmentation specific pathway to control PAX3/PAX7/Prd/Gsbn activity, which would implicate signaling through WNT, FGF and Notch (47). The ability of these signaling cues to operate in a discrete temporal and spatial manner would create regional differences in PAX activity, likely through the control of distinct target gene preferences. Our findings further highlight the complex interplay between the paired domain and homeodomain, as well as the importance of sequences outside of these canonical domains in the control of DNA-binding and functional diversification within the PAX family.

## Acknowledgements

The authors thank N. Hu for technical assistance.

## Author contributions

GNC conducted most of the experiments, analyzed data and contributed to writing of the manuscript. BDS carried out the NMR experiments and their interpretation. DAU conceived the idea for the project, carried out experiments, interpreted data, and wrote the paper with GNC.

## FOOTNOTES

This research was supported by grants from the Alberta Cancer Foundation (grant no. 25675), Alberta Innovates-Health Solutions (grant no. RES20100670), Women and Children’s Hospital Research Institute (grant no. RES0018640 to DAU), and the Canadian Institutes of Health Research (grant no. 37769 to BDS). DAU was supported by the Mary Johnston Chair in Melanoma Research.

The abbreviations used are: PAX, Paired Box; PD, paired domain; HD, homeodomain; EMSA, Electrophoretic Mobility Shift Assay; NMR, nuclear magnetic resonance; 1D, one dimensional

---

**Supplementary Table.**
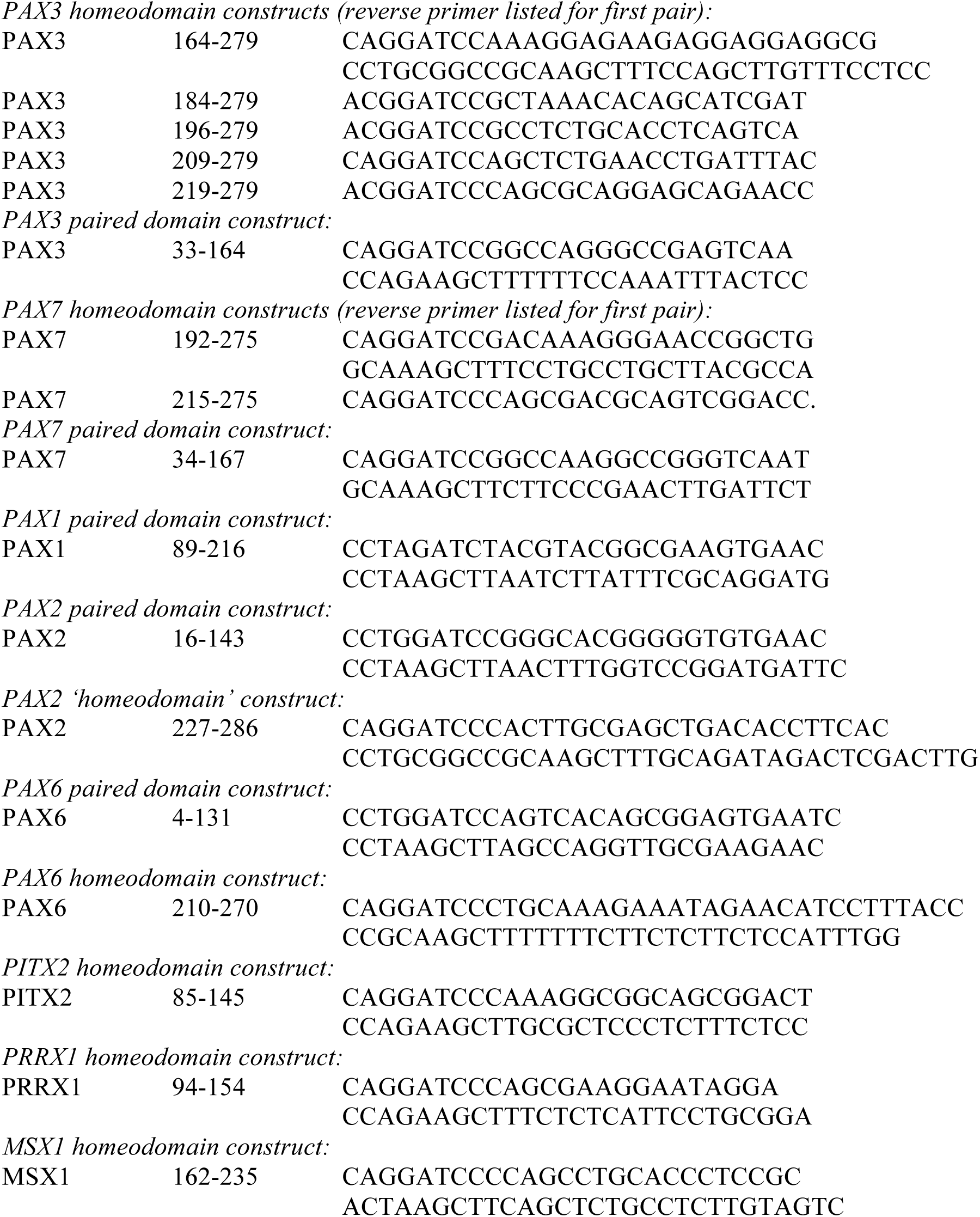
Oligonucleotides used for bacterially expression constructs. Table identifies protein, domain, amino acid position and primer sequences. PAX3 homeodomain constructs (reverse primer listed for first pair):

